# Barrel cortex development lacks a key stage of hyperconnectivity from deep to superficial layers in a rat model of Absence Epilepsy

**DOI:** 10.1101/2022.03.18.484944

**Authors:** Simona Plutino, Emel Laghouati, Guillaume Jarre, Antoine Depaulis, Isabelle Guillemain, Ingrid Bureau

## Abstract

The development of cortical neuronal wiring is precisely orchestrated and goes through several stages, some of which coincide with critical periods when sensory experience is most influential. In particular, although ascending excitatory and inhibitory projections from the deep layer 5 to upper layers are first strong, they recede by the end of the critical period of their targets cells located in upper layers. Alterations in these transient innervations impair the construction of the later circuits that remain at adulthood, but it is unknown whether they could lead to pathologies. Here, we address this question in a genetic model of Absence Epilepsy, a neuro-developmental disease, where epileptogenesis occurs during the postnatal maturation of barrel cortex, the seizure initiation site. Using functional mapping by laser scanning photostimulation with glutamate uncaging in slices, we investigated the pattern of projections onto layers 2/3 pyramidal cells from 2-week old rats. We found that its maturation skipped the key stage during which pyramidal cells received strong projections from both excitatory and inhibitory neurons located in deep layers. At the same age, neuronal activity recorded *in vivo* with two-photon functional imaging was organized in fewer clusters than in control rat pups during this transient hyper-innervation. Later, around the onset of typical absence seizures (∼1 month old), over-excitability of cells was observed across layers. Using this genetic model of childhood epilepsy, we provide first evidence that failure to develop this transient hyper-innervation from deep cortical layers plays a role in pathological neural dysfunctions.

**Significance Statement:** During development of cortex, innervation from deep to upper layers is thought to provide a temporary scaffold for the construction of the circuits that remain at adulthood. Whether an alteration in this sequence causes brain malfunctions in neuro-developmental diseases is unknown. Using functional approaches, we investigated in a genetic model of Absence Epilepsy and control rats the maturation of innervation onto layer 2/3 pyramidal cells of barrel cortex and the cell organization into neuronal assemblies. We found that development in this model lacks this early surge of connectivity with deep layers and the concomitant structuring into multiple assemblies. Later on, at seizure onset, neurons in all layers are hyper-excitable, suggesting this feature of epilepsy develops from prior connectivity defects.

## Introduction

The development of cortical circuits in sensory cortices proceeds from bottom to top layers. In the primary somatosensory cortex of rodents, it begins with the wiring of the cortical plate, goes on during the first postnatal week with the appearance of the so-called barrels, the cell aggregates in layer 4 that receive a topographic thalamic innervations, and it continues with the growth of the projection from layer 4 to layer 2/3. Although ascending projections from layer 5 are weak in the mature barrel cortex (1, 2), it was shown that cells in layers 1 to 4 received strong inputs from excitatory and inhibitory neurons located in layer 5 for a brief period at an early stage of their maturation (3-5). Some of these projections are thought to provide a scaffold to the final wiring that will remain in the adult brain (3). Indeed, the period they are active coincides with the critical period of their target layers, when sensory experience has a great influence (6, 7). Hence, the properties of final circuits in upper layers may strongly depend on these early and transient projections from deep layers. Whether abnormalities in this developmental sequence leads to pathologies remains unknown.

Here we addressed this question in a genetic model of Absence Epilepsy (AE), a developmental form of epilepsy that affects children between 5 and 12 years of age (8). In Genetic Absence Epilepsy Rats from Strasbourg (GAERS), spike and wave discharges (SWDs), the typical EEG pattern of absence seizures, are initiated in the barrel cortex (9-11). We showed that epileptogenesis takes place during cortical development with abnormal oscillations first observed at postnatal day (P) 15 that evolve into *bonafide* SWDs up to P25-30 (12). Simultaneously, the intrinsic excitability of pyramidal neurons increases in deep layers (12). Moreover, spiking responses of neurons evoked by whisker stimulations at P15 are weaker in GAERS despite increased thalamic responses (13). This suggests that abnormalities exist in circuits of barrel cortex at an early stage of development which might set the ground to absence epileptogenesis.

To test this hypothesis, we compared the development and maturation of projections targeting layer 2/3 pyramidal cells in GAERS and control Wistar rats. We used Laser Scanner Photostimulation (LSPS) with glutamate uncaging combined to patch clamp recordings in brain slices to map the functional projections received by L2/3 pyramidal neurons. Experiments were performed during the maturation of L2/3 innervations (∼ P15) and at SWD onset (∼ P30). The structure of early neuronal assemblies of layer 2/3 cells was then investigated with *in vivo* two-photon calcium imaging. Our results support the hypothesis of an altered developmental program in GAERS barrel cortex which deprives layer 2/3 cells from the excitatory and inhibitory drive of deeper layers, especially layer 5, at an early stage. They suggest that abnormalities in the developmental sequence of the intracortical wiring precede and may lead to the enhanced excitability of excitatory and inhibitory neurons in GAERS observed at SWD onset.

## Results

### Evolution of inhibitory and excitatory inputs to L2/3 pyramidal cells in GAERS during cortical development

In order to determine whether cortical development was altered in GAERS, we used LSPS with glutamate uncaging and patch clamp recording (14, 15) in barrel cortex slices (16). Rats were from P11 to P19 (termed “P15” from now on) to cover an active stage of growth and maturation of the projections received by the L2/3 pyramidal cells (14, 17). Glutamate was photo-released on a 16 × 16 pixel grid (90 µm spacing) to excite cells of different layers while a L2/3 pyramidal cell was recorded in voltage-clamp mode (Fig. 1A). Both excitatory and inhibitory inputs across layers were weaker in GAERS rats at P15 than in controls (GAERS EPSC L2/3, - 32%, p = 0.016; L4, - 37 %, p = 0.0028; L5, - 36 %, p = 0.000068; IPSC L2/3, - 38 %, 0.00029; L4, - 58 %, p < 0.00001; L5, - 61 %, p < 0.00001; Fig. 1B,C). However, some of these differences seemed transient as the dynamic of inputs in the Wistar rats showed day-to-day changes in line with previous reports (3, 4, 18), in particular in L4 (excitatory + inhibitory) and in L5 (excitatory) (Fig. 1D; gray symbols). Hence, L4 and L5 inputs increased and then decreased within few days in Wistar rats, whereas they appeared overly stable during the same period in GAERS (Fig. 1D; open black symbols). We repeated these experiments on P29-37 (termed “P30” from now on) rats, at the onset of SWDs. This time, we observed in GAERS an increase across layers in the strength of the projections, both excitatory and inhibitory (Fig. 2A,B). The difference was statistically significant for both projections originating from layers 4 and 5, when compared to Wistar rats (GAERS L4 EPSC, × 2.2, p = 0.00021; L5 EPSC, × 1.5, p = 0.00054; L4 IPSC, × 2.3, p = 0.00020; L5 IPSC, × 2.1, p < 0.00001).

**Figure 1:**
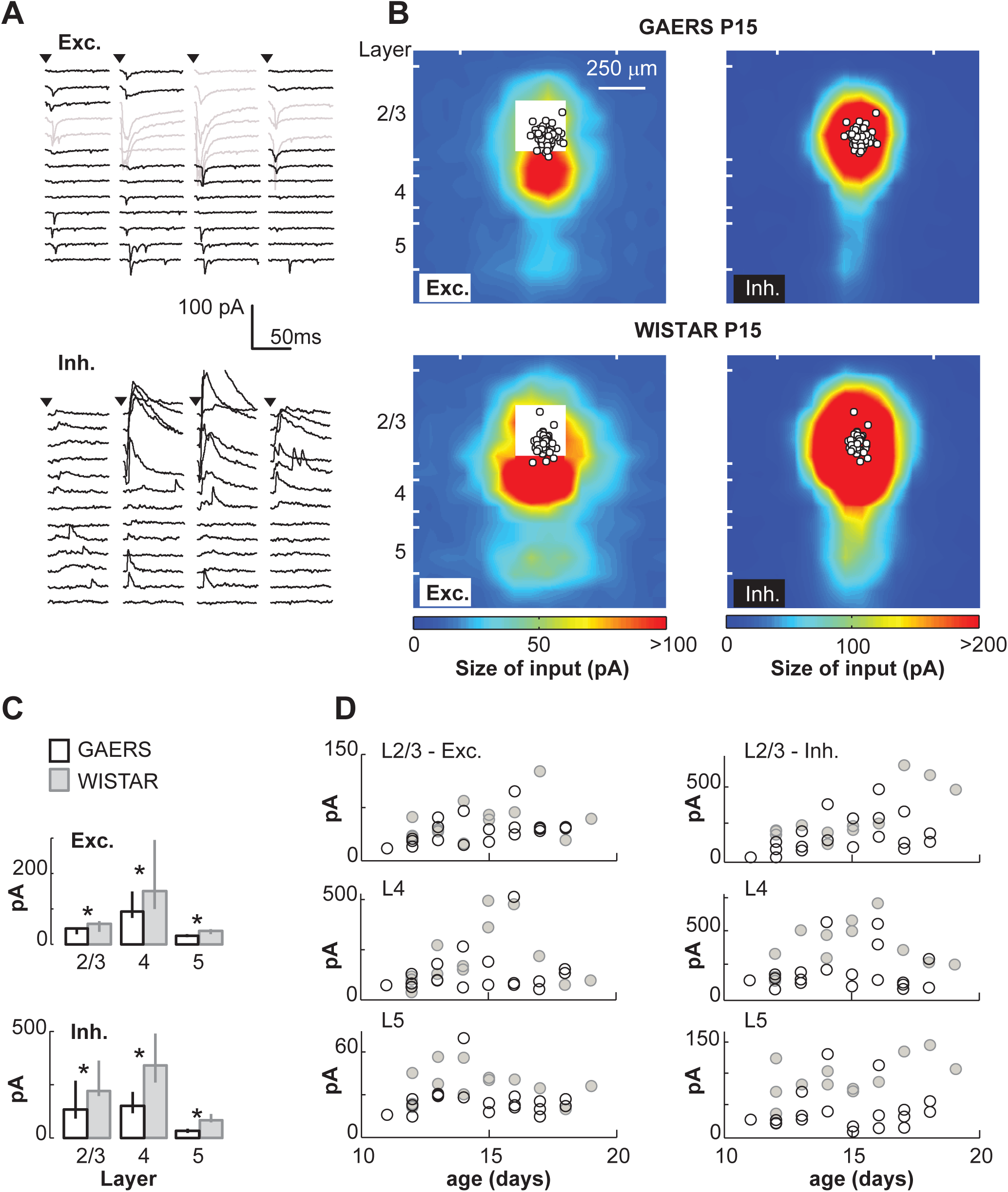
Weaker projections onto layer 2/3 pyramidal cells in P15 GAERS. **A**, Left, Responses evoked by the uncaging of glutamate at cells were recorded in L2/3 pyramidal cells at – 70 mV to record EPSCs (left) and at 0 mV to record IPSCs (right). Each trace is the response at one uncaging site of the LSPS grid. 65 of the 256 traces are shown here. Traces in gray were ignored as they contained direct responses evoked by the uncaging of glutamate onto the recorded cell. One trace of IPSC is cropped. **B**, Averaged maps of excitatory (left) and inhibitory (right) responses recorded in layer 2/3 cells of P15 GAERS (top) and Wistar (bottom). White circles indicate the soma position of the recorded cells in L2/3. Excitatory responses evoked in the white areas close to the recording sites were masked by direct glutamate-evoked responses and could not be analyzed. White ticks on the side of maps indicate the regions of interest for quantification in C. **C**, Bars show the median size and the whiskers the 25-75^th^ percentiles of EPSCs (top) and IPSCs (bottom). **D**, Size of EPSCs (left) and IPSCs (right) as a function of postnatal day. Each circle is a rat (open black : GAERS, solid gray: Wistar). The number of rats and number of cells (N-n) for P15 GAERS EPSCs, 19 – 63; Wistar EPSCs, 13 – 42; GAERS IPSCs, 19 – 64; Wistar IPSCs, 12 – 40.

**Figure 2:**
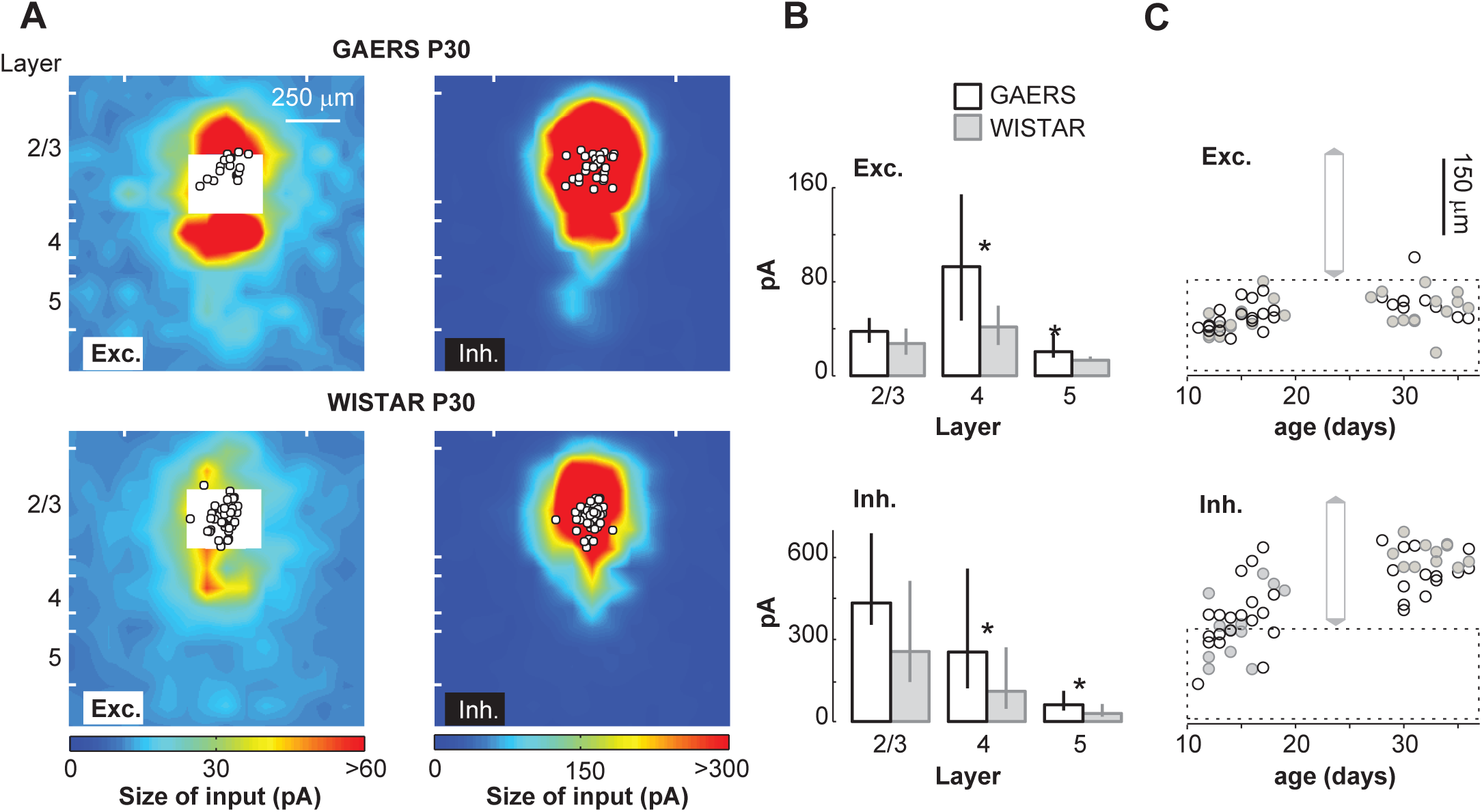
Stronger projections onto layer 2/3 pyramidal cells in P30 GAERS. **A**, same as in Fig.1B for P30 animals. **B**, Bars show the median values and the whiskers the 25-75^th^ percentiles of the size of EPSCs (top) and IPSCs (bottom) evoked by LSPS in L2/3, L4 and L5 in GAERS (open black) and in Wistar (solid gray). **C**, The center of mass of synaptic responses was computed. It shows in Wistars like in GAERS a shift towards the pia, especially when computed from the size of IPSCs. Dashed lines indicate the limits of layer 4. The gray vertical bar indicates the soma position of the recorded cells in L2/3. P30 GAERS EPSCs N-n are 10-16; Wistar EPSCs, 14 - 46; GAERS IPSCs, 14 - 32; Wistar IPSCs, 11 - 44 from A to C. Numbers for P15 GAERS are in Fig.1.

Since GAERS L2/3 pyramidal cells are deprived from inputs at an early stage of their maturation, we asked whether this would alter the timing of the transition to the adult pattern of projections of these cells, as suggested in a recent report (18). We tested this hypothesis by comparing the center of mass of these projections along the maps vertical axis in the two strains (see Methods). In Wistar rats, the center of mass of inhibitory inputs shifted from the middle of L4 up to L2/3 between P11 and P19, illustrating the consequence of synaptic refinements that progressively gives more weight to local GABAergic projections (Fig. 2C). This shift had a similar time course in GAERS. This suggests that the timing for transferring the inhibitory drive of L2/3 cells from deep to superficial layers is preserved in the AE model. Same observation was made for the excitatory inputs. Note that the small shift in the center of mass of EPSCs, from the center to the upper part of L4, may have been underestimated since it was computed from responses evoked at sites that were out of the vertical axis of the recorded cells (see Methods). Altogether, these findings argue against the hypothesis of a developmental delay in GAERS. Rather, they suggest defects in mechanisms controlling the amount of inputs received by cortical cells.

### A hyper-excitability of excitatory and inhibitory cells across layers at SWD onset

Differences in the strength of projection stimulated by glutamate uncaging can result from a difference in (i) the excitation evoked by glutamate, (ii) the connectivity rate and/or (iii) the synaptic strength (19). We investigated the first hypothesis with loose patch recordings to record action potentials (APs) while glutamate was uncaged (Methods; Fig. 3A). Based on their excitation profiles, we calculated the probability at which cells would be activated in the grid used for generating the input maps (see Methods). At P15, this probability was similar between GAERS and Wistar rats, across layers (Fig. 3B). In addition, neurons fired similar numbers of APs once activated at this age (Fig. 3B; L4, p = 0.29; L5, 0.62; L2/3, 0.02, GAERS is × 1.06). In contrast, at P30, the probability to excite neurons in the LSPS grid rate was higher in GAERS, albeit significantly in L5 only (× 1.7; difference between probabilities, 0.37; 95% CI, [-0.23 0.27]). In addition, firing responses of L4 neurons had more APs in GAERS than in Wistar rats (p = 0.029; Fig. 3B). Furthermore, consistent with a higher excitability of the excitatory cells in the slice at P30, EPSCs evoked by LSPS in L2/3 and L4 had shorter latencies in GAERS at this age (L2/3, - 2 ms, p = 0.000053; L4, - 3 ms, p < 0.00001; Fig 3C) whereas latencies were similar at P15 (Fig. 3C; L2/3, p = 0.39; L4, 0.03; L5, 0.17). Current-clamp recordings confirmed that neurons were hyper-excitable at P30. Indeed, layer 5 pyramidal cells were more depolarized at rest in GAERS than in Wistar rats (+ 2 mV; p = 0.025; Fig. 3D), in line with our previous findings *in vivo* (12). Layer 4 stellate cells were also more depolarized at rest in GAERS (+ 1.5 mV; p = 0.028), whereas we found no significant difference for layer 2/3 pyramidal cells (p = 0.75). We then compared for each neuronal subtype the number of APs evoked by increasing steps of depolarization from a holding potential of -70 mV, to eliminate differences due to the fact that GAERS cells were depolarized at rest. As previously observed *in vivo* (12), we found no difference in the firing responses of pyramidal cells in layer 5 and 2/3 between P30 GAERS and Wistar rats in this condition (Fig. 3E). However, L4 cells in GAERS had an increased functional gain: 67 % fired at 100 pA vs 24 % in Wistar, and firing remained stronger in GAERS up to the 250 pA step (> 95% confidence intervals; Fig. 3E). To summarize, we found that at P15 excitatory neurons across layers fired at similar levels in GAERS and Wistar rats, suggesting that other defects accounted for the weaker synaptic inputs received by L2/3 pyramidal cells in the AE model. However, the stronger inputs these cells received at P30 is likely explained by a greater excitability of excitatory cells of deeper layers.

**Figure 3:**
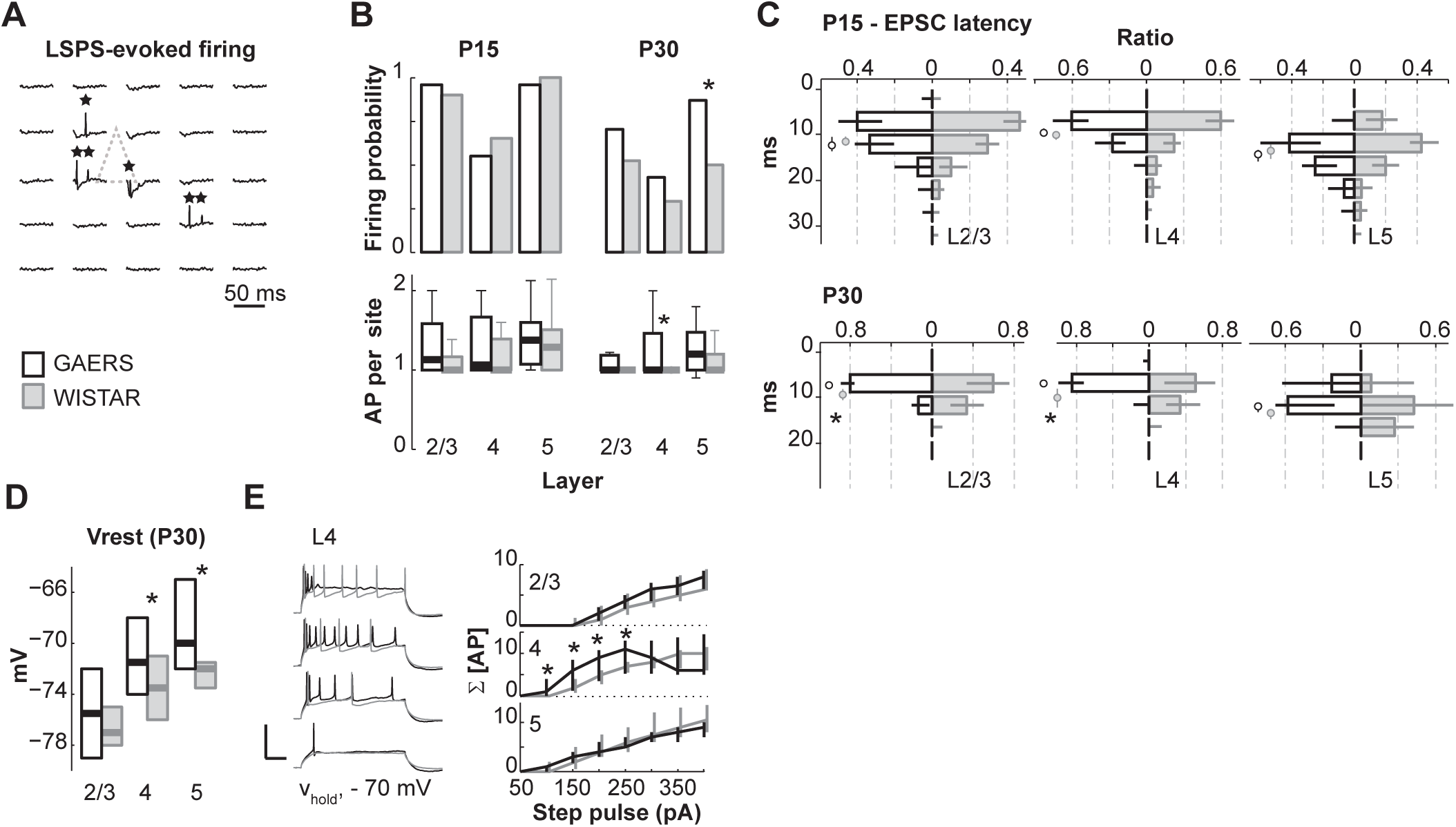
GAERS excitatory cells have abnormal excitability at P30, but not at P15. **A**, Glutamate was uncaged on a 8 × 8 – 50 μm spacing grid while APs from single cells were recorded in the loose patch mode. 25 of the 64 traces from a GAERS L5 cell are shown here. Stars indicate APs, the dashed line the position of the soma in the grid of stimulations. **B**, Top, the probabilities (top) and the mean number of spikes (bottom) that cells fired upon LSPS in the 90 μm spacing grid used for generating synaptic input maps. In the bottom plot, thick lines are the median values, the bars the 25-75^th^ percentiles, the whiskers contain the values within the percentiles × 1.5. P15 GAERS N = 7, n = 24 – 20 – 24; P15 Wistar, N = 8, n = 30 – 23 – 21; P30 GAERS, N = 8, n = 27 – 28 – 23; P30 Wistar, N = 10, n = 21 – 24 – 18. **C**, Distributions in GAERS (open black) and Wistar (solid gray) of the latencies to the onset of EPSCs evoked in the synaptic input maps (5 ms bins) for P15 (top) and P30 animals (bottom). Circles and whiskers are the median values and 25-75^th^ percentiles. Same N-n as in Fig.1 and 2. **D**, L2-5 excitatory cells were recorded in current clamp in P30 animals. The thick lines are the medians and the bars the 25-75^th^ percentiles of their membrane potential at rest (Vrest) in GAERS (open black) and in Wistar (solid gray). **E**, Example traces (left; layer 4) and medians (right) of cell firing responses evoked by steps of increasing currents (100 – 250 pA, bottom to top) in P30 GAERS (black) and Wistar (gray) rats. Cell membrane potential was held at - 70 mV in this experiment. Scales bars, 70 mV and 100 ms. GAERS, N = 6, n = 14 – 18 – 14 (L2/3 – L4 – L5); Wistar, N = 7, n = 15 – 20 – 12.

Two observations suggest that the firing properties of interneurons undergo similar alterations than for excitatory neurons in GAERS during barrel cortex development. First, we observed no difference in the latencies of IPSCs in GAERS and Wistar rats at P15 (Fig. 4A; L2/3, p = 0.22; L4, 0.93; L5, 0.044) whereas these were shorter in GAERS at P30 (L2/3, - 4 ms; p = 0.000012; L4, - 3 ms, p = 0.0022; L5, - 6 ms, p = 0.000015). This suggests that, alike excitatory cells, GABAergic interneurons across layers become more excitable in the AE model at P30. We sought to further explore this finding. Indeed, the long tail in the histograms of latencies of IPSC evoked by stimulations in L5 and L2/3 suggested that two populations of interneurons, with either quick or delayed excitation, innervated the L2/3 pyramidal cells (Fig. 4A-C). In contrast, a tighter distribution of IPSC latencies suggested that innervation originating in L4 was largely from a single population with a quick excitation (Fig. 4A,B). This is consistent with a previous report that fast-spiking (FS) interneurons fire APs upon glutamate uncaging with shorter latencies than other GABAergic interneurons in barrel cortex (20). At P30, short-latency IPSCs (sl-IPSCs; latency < 15 ms) had larger amplitudes than long-latencies IPSCs (ll-IPSCs) in both GAERS and Wistar rats (Fig. 4C,D). We hypothesized that the firing of over-excitable neurons should resist better to a decrease in stimulation intensity. Therefore, in a subset of neurons, we lowered the laser intensity to decrease the concentration of uncaged glutamate and the depolarization of interneurons. The stimulation grid was kept the same, permitting to analyse the change at every site, separately for sl-IPSCs and ll-IPSCs. Data of L2-5 were pooled. Decreasing the stimulation intensity resulted in a moderate loss of evoked sl-IPSCs in the grid in P30 GAERS, ∼ - 30 %, which was significantly lower than in Wistar (- 65 %; p = 0.00029 Wilcoxon; Fig. 4C,E). This loss was similar for ll-IPSC in the two strains. This is consistent with a pronounced excitability of FS interneurons in the AE model. Altogether, our findings suggest that the intrinsic properties of excitatory neurons and fast-spiking interneurons in GAERS are changed in parallel, with an abnormal increased excitability emerging between P20 and P30.

**Figure 4:**
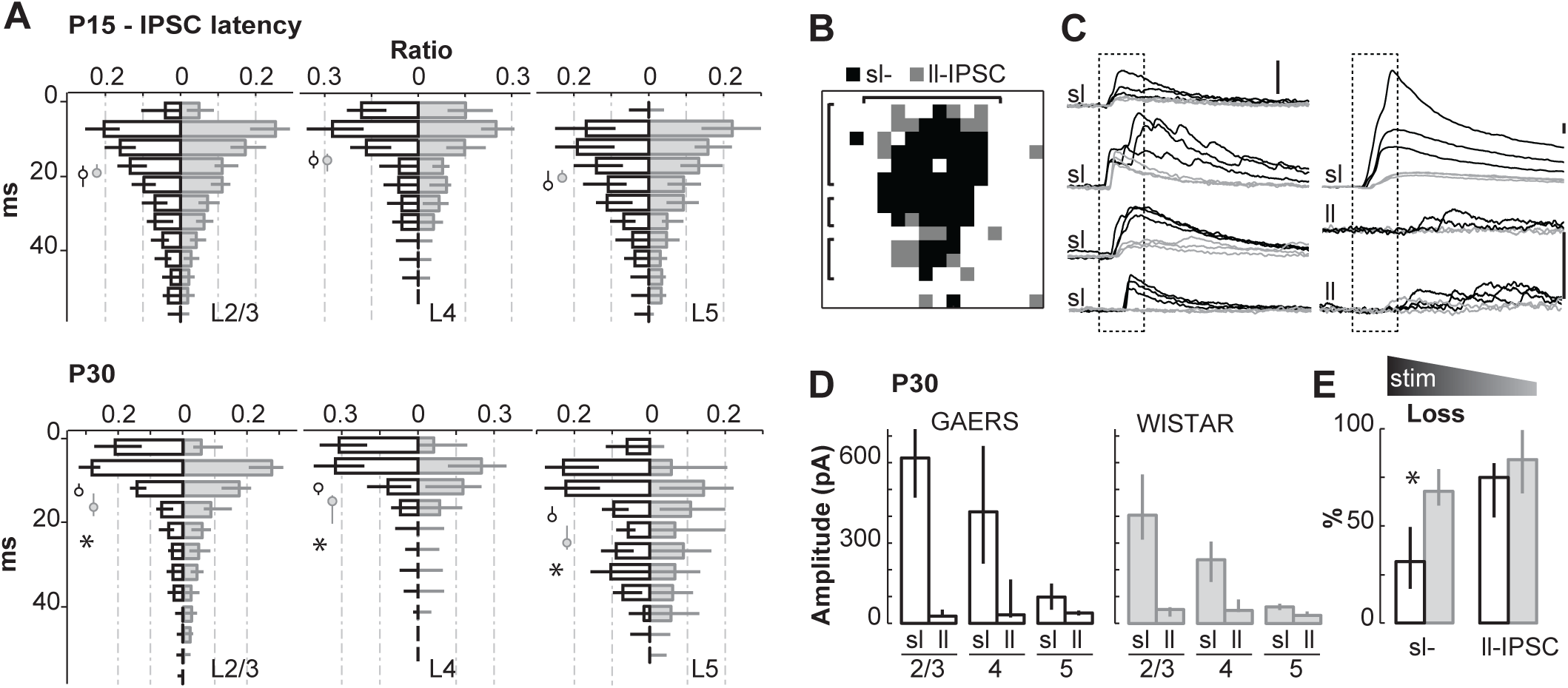
GAERS inhibitory cells are hyper-excitable at P30. **A**, Distributions in GAERS (open black) and Wistar (solid gray) of the latencies to the onset of IPSCs (5 ms bins) for P15 (top) and P30 animals (bottom). Circles and whiskers are the median values and 25-75^th^ percentiles. **B**, Example map showing the positions of short latency (sl; < 15 ms) IPSCs (black pixels) and long latency (ll; ≥ 15 ms) IPSCs (gray pixels) of a GAERS cell. Brackets indicate the L2/3, L4 and L5 regions of analysis shown in D. **C**, Example traces of sl-IPSCs and ll-IPSCs in a GAERS cell. Each subpanel is for a single site. The dashed boxes mark a 15 ms time window starting at the stimulus onset. Scale bars are 150 pA everywhere. The nAmp-large sl-IPSCs on the top right were evoked at a stimulation site near the soma of the recorded cell. In black, the responses evoked in control condition (20 mW laser) and in gray the responses evoked with a weaker stimulation (5 mW, see panel E). **D**, Peak amplitude of sl-IPSCs and ll-IPSCs in GAERS (open black) and Wistar (solid gray). **E**, Percentage of lost sl-IPSCs and ll-IPSCs after decreasing the laser power from 20 to 5 mW to lower the concentration of uncaged glutamate. Data of L2-5 were pooled. See panel C for examples. In panels A-D, same N-n as in Fig. 1 and 2. For panel E, GAERS N-n are 4 – 11; Wistar, 6 – 16.

### Hypo-connectivity during the second postnatal week in GAERS

Our data indicate that the globally weak input received by GAERS L2/3 cells at P15 is not due to abnormally weak excitability of inhibitory and excitatory cells. We then explored the alternative hypothesis of a decreased connectivity in GAERS. Given that cells fired at similar levels in GAERS and Wistar rats at P15 (Fig. 3,4), differences in the rate with which synaptic responses were evoked in the LSPS grid would indicate differences in connectivity. This one is proportional to the density of cells under the grid and to the probability that each cell is connected to the recorded neurons in layer 2/3 (19). Therefore, we computed maps of response rates (Fig. 5A) and observed a lower probability of evoking EPSCs and IPSCs in P15 GAERS when compared to Wistar rats. Differences were statistically significant across layers, although they seemed largest for stimulations in L5 (L5 EPSC, - 52 %, p < 0.00001; L5 IPSC, - 29 %, p < 0.00001; L4 EPSCs, - 29 %, p < 0.00001; L4 IPSCs, - 17 %; L2/3 EPSC, - 19 %, p = 0.0040; L2/3 IPSCs, - 19 %, p = 0.00028; Fig. 5B,C). Maps of response probabilities showed a significant decrease in the rate of sl-IPSCs too (L5, - 40 %, p < 0.00001; L4, - 17 %, p = 0.00001; L2/3, - 20 %, p = 0.00028; ll-IPSC, p = 0.70 - 0.26 - 0.10; Fig. 5C). The connectivity level was regulated during this developmental period in Wistar rats (4), in particular in layers 4 and 5 as the rate of EPSCs increased to reach a peak around P15 and decreased thereafter (Fig. 5D). On the contrary, these rates were stable in GAERS during the same period. However, the pattern of projections onto L2/3 cells still underwent changes in this strain. In particular, the rate of sl-IPSCs evoked by the stimulation of L2/3 interneurons increased in GAERS the same way as in Wistar rats (Fig. 5D). Furthermore, the center of mass of excitatory and inhibitory responses computed for each cell moved upward in the LSPS grid between P11 and P19 in GAERS like in Wistar rats (Fig. 5E), which is consistent with a change in the pattern of connectivity.

**Figure 5.**
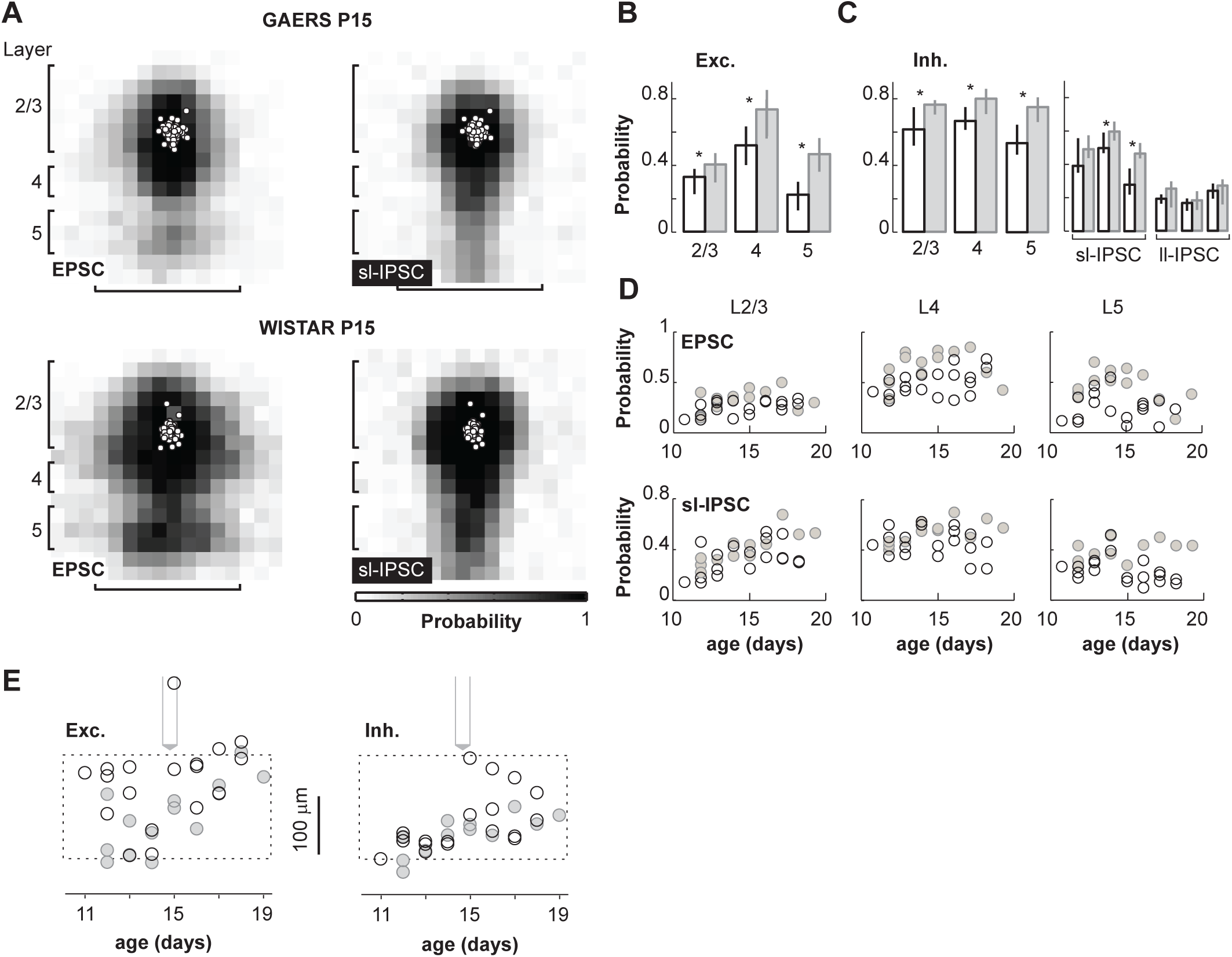
Functional connectivity is lower in P15 GAERS. **A**, Maps of response probabilities for EPSCs (left) and IPSCs (right) in P15 GAERS (top) and Wistar (bottom). Black vertical and horizontal brackets are the regions of interest for quantifications in B and C. **B**, Probabilities of evoking EPSCs computed by layer. **C**, Same for IPSCs, sl-IPSCs and ll-IPSCs. **D**, The probability of responses as a function of postnatal days, for EPSCs (top) and sl-IPSCs (bottom). Each symbol is a rat. **E**, The center of mass computed from the probabilities of EPSCs (left) and IPSCs (right), independently of their amplitude. Same representation as in Fig. 2C. Same N-n as in Fig. 1.

Finally, we explored the third hypothesis that a diminished synaptic strength could also explain the weaker projections in GAERS at P15. This requires to stimulate pre-synaptic cells singly. The probability to evoke synaptic responses was not homogeneous across sites of the LSPS grid as expected from the organization of the projections targeting L2/3 cells. We thus hypothesized that stimulations at sites with low success rates could evoke unitary responses due to the fact that only one cell was connected to the recorded neuron. We found that EPSCs and sl-IPSCs evoked at sites with probabilities < 0.1 had similar amplitudes in GAERS and Wistar rats (median [25^th^ 75^th^ percentiles]: GAERS EPSC, 28 [25 3225 32] pA; Wistar, 30 [25 41] pA; p = 0.58; GAERS IPSC, 45 [35 60] pA; Wistar, 71 [40 91] pA; p = 0.13, Fig. 6). This suggests that the weak inputs received by the GAERS L2/3 cells at P15 are not due to a decrease in synaptic strength.

**Figure 6.**
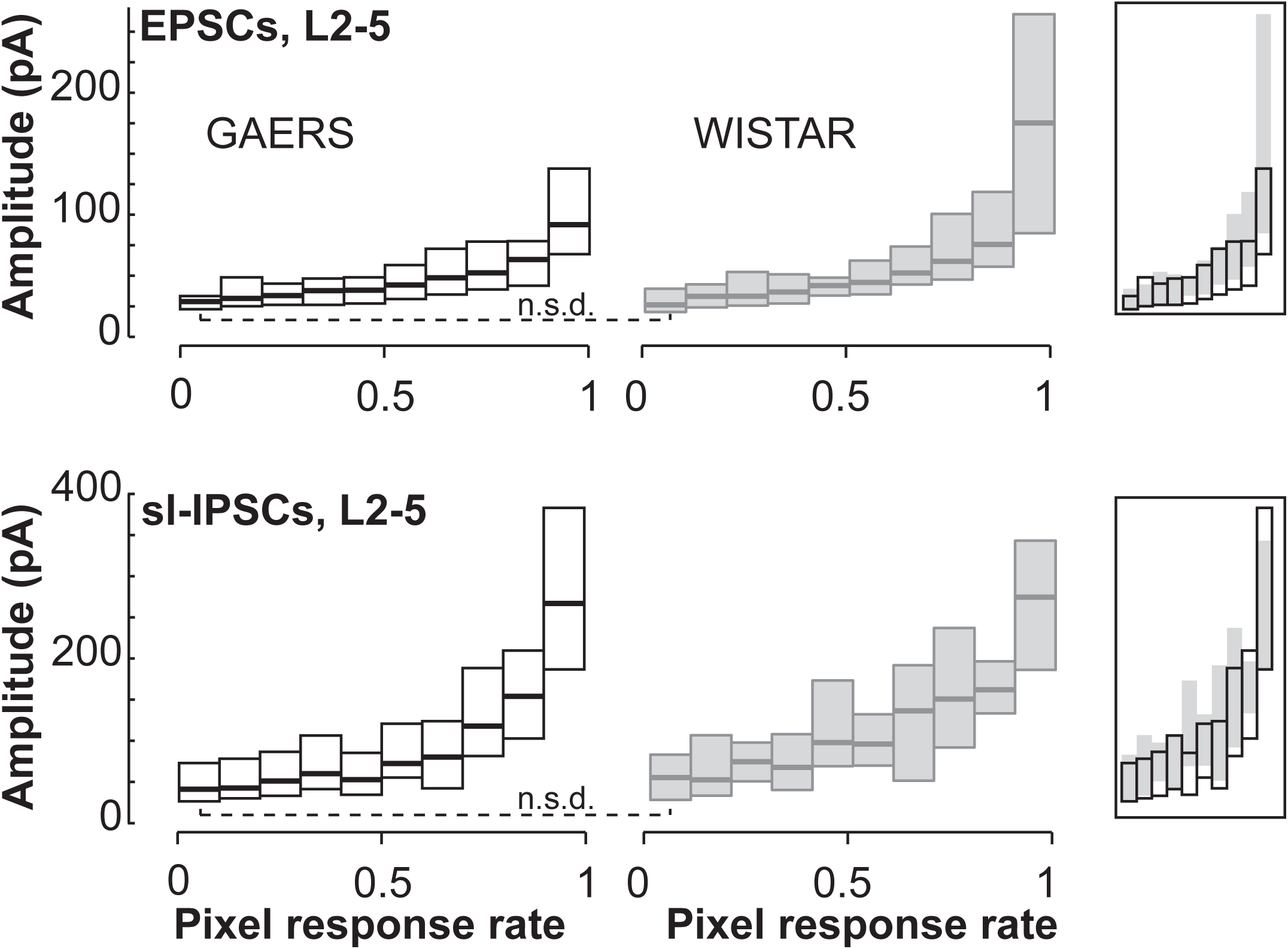
The synaptic strength of excitatory and inhibitory connections is similar in P15 GAERS and Wistar rats. Amplitude of EPSCs (top) and sl-IPSC (bottom) as a function of the probability of responses at the stimulation sites. Thick lines are the median values, the bars the 25-75^th^ percentiles. Response amplitudes were not significantly different (n.s.d.) in Wistar rats and GAERS at sites with rates < 10 %, which suggests synaptic strength of single connections were similar in the two strains. In insets on the right, the overlays. Same N-n as in Fig. 1.

### An abnormal clustering of L2/3 neurons coincides with the hypo-connectivity at P15

The connectivity defects in GAERS observed at P15 should have consequences on the way neurons are recruited during the activation of the barrel cortex. Spontaneous activity at the end of the first postnatal week is characterized by bursts of coordinated activity that recruit large numbers of cells in superficial layers of sensory cortices (21). This early pattern evolves into a de-correlated state in which activity is continuous and neuronal assemblies with synchronized firing are small. It was shown that in the primary auditory cortex, this so-called de-correlation process (22) was transiently paused at the same time superficial cells were innervated by cells in deep layers (4). This innervation from deep layers was low in P15 GAERS, therefore we hypothesized that abnormalities in the maturation of neuronal assemblies should be observed. The spontaneous firing activity of layers 2/3 neurons was monitored using *in vivo* two-photon calcium imaging (Fig. 7A-C). About similar high number of active cells contributed to the global activity of layer 2/3 in GAERS and Wistar rats (median ratio of active cells over all cells [25^th^ 75^th^ percentiles]: GAERS, 0.90, [0.88 .96]; Wistar, 0.88 [0.81 0.92], p = 0.005, × 1.03 in GAERS). However, GAERS neurons were ∼ 2 times more active (p < 0.00001; Fig. 7C, D). This may be caused by the low inhibitory innervation these cells receive at this stage. To characterize the structure of active neuronal circuits, we identified pairs of neurons whose activity was significantly correlated over the course of the imaging session (Methods). As expected, the fraction of correlated pairs was low in Wistar rats (< 0.5), confirming that the de-correlation process had been initiated (Fig. 7E) (21). This ratio was ∼ 40 % higher in GAERS (Fig. 7E, GAERS: p = 0.0038), whereas the correlation coefficient of those pairs was lower (Fig. 7F, median [25^th^ 75^th^ percentiles]: GAERS, 0.10 [0.10 0.12]; Wistar, 0.14 [0.13 0.18]; p < 0.00001). Neurons were then grouped into functional clusters by applying an algorithm that sequentially joins cells that have statistically significant similarities in their firing pattern (Methods; Fig. 7C) (23, 24). Clusters were 2.5 times less numerous in GAERS (1.0 [1.0 2.1]; Wistar, 2.6 [2.1 2.7]; p = 0.0313). In addition, we found a single cluster in 64 % of recordings in GAERS vs 20 % in Wistar rats. (Fig. 7H). This difference was observed at P14-P16 after which (P17-P19) the number of clusters decreased in Wistar rats, concomitantly with the decrease in the strength of projections from deep layers that we observed in LSPS maps (Fig.7H). On average, clusters had more cells in GAERS than in Wistar rats (GAERS, 17 [7.5 28]; Wistar, 5 [4.1 8.6]; p < 0.00001; Fig. 7I) but cell densities were similar in both strains (µm^2^/neuron; GAERS, 693 [454 810]; Wistar, 747 [705 925]; p = 0.1346) (Fig. 7J). These findings indicate that the developmental sequence organizing L2/3 neurons of barrel cortex into successive networks is disturbed in GAERS since cells were integrated in few large functional clusters at a time they should have been distributed in several smaller ones.

**Figure 7.**
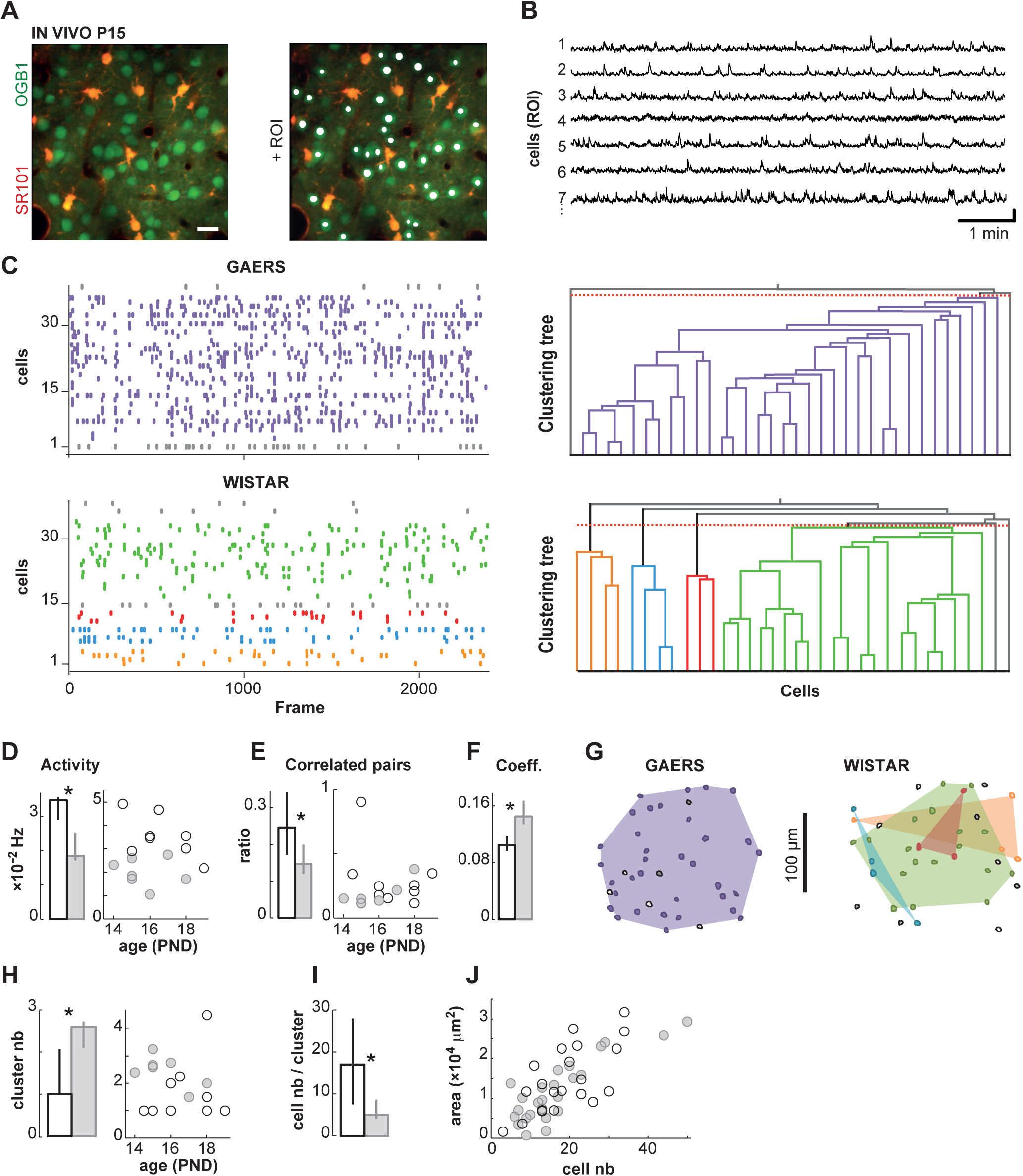
Altered clustering of neuronal activity in P15 GAERS barrel cortex *in vivo*. **A**, Left, typical field of view obtained after multicell bolus loading with the calcium indicator OGB1 (green) and the glial marker SR101 (red) in the layer 2/3. Right, regions of interest (ROI) are shown in white overlaying neuron somata. Scale bar: 20 µm **B**, Examples of neuronal calcium activities. Scale bars: 50 % and 1 min. **C**, Example raster plots of calcium events (left) and corresponding clustering trees (right) in P15 GAERS and Wistar rats. Each color corresponds to a cluster. In the clustering trees, the red dashed lines indicate the statistical significance cut-off. **D**, Bars, frequency of calcium events during the recording session. Whiskers are 25-75^th^ percentiles. Symbols, frequency as a function of postnatal days. Each symbol is a rat. GAERS, black open; Wistar, solid gray. **E**, Ratio of significantly correlated pairs over all possible pairs of active cells. **F**, Coefficient of correlation for significantly correlated pairs of cells. **G**, Spatial representation of the clusters shown in C. Unactive cells (gray) and unclustered cells (white) are shown. **H**, Number of clusters per recording. Median (bars) and values as a function of postnatal days (symbols). **I**, number of cells in clusters. **J**, Area of clusters as a function of the cell number inside. Only the largest cluster of each recording is shown. GAERS N-n are 9 – 22; Wistar, 7 – 25.

## Discussion

We investigated the properties of neurons and networks in the barrel cortex of Wistar rats from P11 to P36 and compared them with those in a model of AE. During the second postnatal week, a surge of connectivity occurred in the Wistar rats which transiently connected excitatory and inhibitory neurons in L5 to the pyramidal cells of L2/3 at the same time neuronal activity in L2/3 was structured in multiple assemblies. This phenomenon did not occur in GAERS where neuronal connectivity remained weak and the activity structured in one or few assemblies in L2/3 during this period. At P30, around SWD onset, excitatory and inhibitory neurons of barrel cortex were hyper-excitable and projections from all layers were stronger onto the cells of L2/3 in GAERS.

### A lack of projections from deep layers in P15 GAERS

The maturation of cortical circuits goes through different stages with sometimes abrupt transitions. Transient patterns of synaptic connections were repeatedly observed in the developing cortical plate: they concerned all neuronal subtypes, glutamatergic as well as GABAergic connections (3, 4, 18, 25). Some patterns are thought to regulate the early cortical activity of thalamic origin. Hence, GABAergic interneurons in layer 5 innervated by the thalamus connect layer 4 stellate cells ∼ P5 (3, 18) and layer 2/3 pyramidal cells ∼ P9 (3, 4) before receding by the end of the critical period of each layer. Early projections are critical for the maturation of pending circuits and preventing the early GABAergic signalling from layer 5 was shown to delay the formation of adult-like connections (18). Here we report that, in GAERS, connectivity to layer 2/3 pyramidal cells is low around P15 when compared to Wistar rats. However, this appears different from a developmental arrest or delay since refinements in the pattern of projections are still visible in GAERS. Indeed, the center of mass of excitatory and inhibitory inputs received by each L2/3 pyramidal cells moved upwards in the direction of the cell home-layer during this period (Fig. 2C and Fig.5E). In Wistar, this move was driven by the weakening of projections from deep layers, together with the establishment of new connections from superficial layers (4, 18). In GAERS, it was mostly driven by the addition of connections from upper layers since connectivity from deep layers was already low. *In vivo*, neuronal activity in GAERS L2/3 at P15, although stronger, was organized into fewer clusters than in Wistar rats. In these control rats, cluster number diminished after P16 concomitantly with the weakening of projection from deep layers, in line with a previous report (4). We did not observe this sequence in GAERS, which corroborates the hypothesis that a transient increase of inputs from deep layers and a more structured activity in layer 2/3 are linked (4).

Could an abnormal activity be at the origin of this developmental alteration in GAERS? Sensory deprivations impair the transition from a sub-granular to an intralaminar bias in the GABAergic innervation onto cells in layer 4 (18). In contrast, our data suggest that the stage of profuse connections from deep layers is lacking in GAERS. We reported previously that the firing response of thalamic cells evoked by whisker stimulation was larger in P15 GAERS as compared to Wistar rats (13). These findings argue against the hypothesis of a weak thalamic drive. Genetic manipulations that either hastened the formation of local connections between parvalbumin (PV) interneurons and layer 4 stellate cells (Neuregulin-1-ErB4 over-signalling), or blocked the synaptic transmission from distant somatostatin interneurons located in L5 (SNAP25 conditional knock-out) (18), lead to a profound decrease in connectivity with interneurons of deep layers. However, they also provoked a commensurate increase in the strength of local projection from PV interneurons which we did not observe in GAERS pups. Investigating the activity and thalamic wiring of interneurons located in deep layers could shed light on the origin of the defect in connectivity that targets upper layer cells. Possible mechanisms would regulate neuronal density in cortex, such as neurogenesis and migration, as well as the innervation of their target cells (26). The defects we observed in GAERS inhibitory projections appear to concern FS interneurons principally which, likely, expressed PV. It was shown that PV interneurons formed neuronal assemblies of smaller size compared to that made of somatostatin interneurons (27). Our *in vivo* observation that L2/3 neuron activities in GAERS often clustered in a single large assembly, while they formed several smaller clusters in Wistar rats at the same age, also points to a determinant role of PV interneurons in this process. However, differences in the inputs and/or in the connectivity from somatostatin interneurons may have been overlooked as responses from non-FS interneurons were under-represented in the synaptic input maps in our study. Indeed, ll-IPSCs that may have been evoked at the same sites in the LSPS grid than sl-IPSCs could not be measured.

### A generalized increase in neuronal excitability in GAERS barrel cortex follows a transient connectivity defect

Our previous study described the progressive changes in L5 pyramidal cells excitability in GAERS that preceded SWD onset (12). Here, our findings suggest that this alteration may in fact be generalized to all neurons in the barrel cortex at P30, albeit with a maximum in deep layers. We collected direct evidences concerning glutamatergic cells as we saw in all layers a shift of their resting potential toward depolarized values that was significant in L4 and L5, and a tendency in L2/3. Two observations also point to a simultaneous hyper-excitability of the GABAergic interneurons in GAERS: IPSC latencies to onset were shorter and evoked sl-IPSCs resisted better to a reduction of stimulation intensity. The inverse correlation between the excitability of neurons and the amount of inputs they receive is known (28, 29) and a hyper-excitability in GAERS could indicate that the connectivity alteration persisted at P30. However, several findings argue against this. First, the increase in the excitation level of L5 excitatory cells (× 1.7 in GAERS) well accounted for the increase in EPSCs that were evoked with LSPS (× 1.5). Second, the connectivity rate in the excitatory projections from L4 and L5 changed over a few days in Wistar rats, and the differences with GAERS had mostly disappeared by P20. In addition, our recent investigations, based on viral tracing of neurons that are pre-synaptic to layer 2/3 pyramidal cells, found that connectivity was enhanced in 2-3 month old GAERS (30). This increase could be an evolution of the pathology and/or an outcome of the recurrence of SWDs.

Hence, we propose that the cell hyper-excitability at P30 is rather a consequence of dysfunctions in networks that were active around P15. A sequence with an early hypo-connectivity (19), concurrent with an altered de-correlation process of neuronal activity (31), and a late cell hyper-excitability (32, 33) was observed in the cortex of a model of Fragile X syndrome, another neuro-developmental disorder plagued with epilepsy (34). Our present findings suggest that developmental abnormalities which deprive cells of inputs and alter their assembly in networks at a critical time may lead to the hyper-excitability of cells, epilepsy and potentially abnormal brain functions.

## Materials and methods

### Animals and ethics

We used GAERS pups (11–36 days old) of the Grenoble colony and age-matched control Wistar rats (Charles River) of either sex. Animal procedures fulfilled both ARRIVE (Animal Research: Reporting of In Vivo Experiments) and Basel declaration (http://www.basel-declaration.org) guidelines, including the 3R concept. Protocols were approved by our local ethical committees and the French Ministry of Research (APAFIS#9761-201712514393900 v3). Experiments were carried out in accordance with European Union guidelines (Directive 2010/63/EU).

### Brain slices preparation and electrophysiology

Animals were anesthetized with isoflurane (4 % in 1 % oxygen) prior decapitation. Solutions and procedures are in (35).

### Laser Scanning PhotoStimulation with glutamate uncaging

LSPS recirculating solution was complemented with (in mM): 0.2 MNI-caged glutamate (Tocris, Bristol, UK), 0.005 CPP [()-3-(2-carboxypiperazin-4-yl) propyl-1-phosphonic acid], 4 CaCl_2_, and 4 MgCl_2_ (Sigma Aldrich). Traces of whole-cell voltage-clamp recordings were sampled and filtered at 10 kHz. Focal photolysis of caged glutamate was accomplished with a 2 ms 20 mW pulse of a UV (355 nm) laser (DPSS Lasers Inc.) through a 0.16 NA 4 × objective (Olympus). 5 mW pulses were used in a subset of experiments. The stimulus pattern for LSPS mapping of synaptic inputs (i.e., EPSCs or IPSCs) was a 16 × 16 grid (spacing, 90 μm) aligned horizontally with the L4/L5A boundary and vertically with the center of the barrel below the cell. The laser was moved in a spatial pattern designed to avoid consecutive glutamate uncaging over neighboring sites (15). LSPS was also used to quantify the generation of spikes in neurons recorded in loose-patch. Then, the LSPS grid was smaller and centered on soma (8 × 8 grid; spacing, 50 μm).

### Analysis of LSPS data

Synaptic input maps for individual neurons were constructed by measuring the peak amplitude of EPSCs evoked in a 50 ms time window, 5 - 7 ms after the UV stimulus for each position of photostimulation. The time window was 70 ms, 2 ms after the uv stimulus for IPSCs. Typically, two to three maps were obtained per cell and averaged. Direct responses were evoked at sites close to recorded cell-body and EPSCs evoked at these sites were not analyzed. Averaged single-cell maps were used to compute group-averaged maps. Interpolation was performed on averaged synaptic input maps for display purposes only. The center of mass of inhibitory responses was computed from values in the central 4-pixel-column (270 μm width) of the LSPS grid (Fig. 2C, bottom panel; Fig. 5E, right panel). Because direct responses masked large EPSCs evoked at some sites in L2/3, the center of mass of excitatory responses was computed from pixel values in two 2-pixel-columns off the vertical axis of the recorded cells (Fig. 2C, top panel; Fig. 5E, left panel). Computation was as follow: Σ (α × distance from the L4/L5A boundary) / Σ (α) where α was the peak amplitude of EPSCs or IPSCs (Fig.2C) or the occurrence of a synaptic response (Fig.5E). Traces from loose patch recordings were analyzed for counting the number of APs evoked in a 50 ms time window immediately after the stimulus to compute the total number of APs elicited in the grid (nAP_50_) and the mean number of APs per site. At P30, nAP_50_ of L5 cells was larger in GAERS: Wistar, 3.5 [1 5]; GAERS, 6 [4.6 9.8] (median [25^th^ 75^th^ percentiles]; p = 0.00021). No significant difference was found for the other layers and at P15: P30 L2/3, Wistar, 4 [2 6]; GAERS, 5 [3 8.8]. P30 L4, Wistar, 2 [1 4]; GAERS, 2.5 [1 6]. P15 L5, Wistar, 15 [9 22] ; GAERS, 13 [9.5 17]. P15 L2/3, Wistar, 11 [8 14]; GAERS, 10.5 [7.5 22]. P15 L4, Wistar, 5 [3 9.8]; GAERS, 5 [2 10]. The excitation profiles were obtained with an uncaging grid with a 50 μm spacing. To obtain the rate at which cells would be efficiently excited in the 90 μm spacing grid used for generating synaptic input maps, we first computed for each cell nAP_90_ as follow: nAP_90_ = nAP_50_ × (50/90)^2^. Then, the projected rate of excitation was the ratio of cells with nAP_90_ ≥ 1.

### Surgery for calcium imaging

Rat pups were anesthetized with isoflurane (1-3%). A craniotomy was performed over the left somatosensory cortex (AP: –1.2 mm ML: + 4.8 mm from bregma) and the dura matter was removed. Multicell bolus loading of Oregon-Green BAPTA 1-AM (OGB-1, Life Technology) was performed (36) and sulforhodamine 101 (100 μM, Sigma) was added to the solution to identify glial cells (21). The injection was performed at 150-200 μm below the brain surface. The craniotomy was then filled with agarose (1.2% in ACSF, type III-A, Sigma) and sealed with a glass coverslip. Recordings started > 1 hour later (36). ECoG monopolar electrodes (enameled copper wire, 220 μm, Block Germany) inserted over frontal and parietal cortices and cerebellum and a computer-based system (PlusEvolution, Micromed, France) enabled selecting periods without abnormal oscillations as these may change our measurements (12). Analysis were conducted using System PlusEvolution, Micromed, France. Rat pups were maintained in a sedated and analgised state by repeated injections of sufentanyl (3 μg/kg (30 min)^-1^, i.p, Jansen) and haloperidol (60 μg/kg (30 min)^-1^, i.p, Jansen) during the recording session (12). At the end of the experiment, animals were euthanized and histology ensured the position of the craniotomy.

### *In vivo* two-photon calcium imaging

We used a LSM 7 MP (Zeiss, Germany) equipped with a 20 × water-immersion objective lens (NA 1.0; Zeiss). The power of Ti:sapphire laser (940 - 950 nm; Chameleon vision II; Coherent, UK) was ≤ 60 mW at the objective. A dichroic mirror (562 nm, Semrock, USA) separated the SR101 and OGB1 fluorescence, detected by two non-descanned photomultiplier tubes (Semrock, USA) with 593/46 nm and 525/50 nm filters. Fields of view were scanned between 3-4 Hz. Before recordings, a z-stack of the loaded area was acquired to help post-hoc analysis (i.e. to ensure that the focal plane was located near the neurons equator (21). The imaged fields were 301 [226 410] ×10^2^ µm^2^ in GAERS (N = 9; n = 22) and 346 [306 376] ×10^2^ µm^2^ in Wistar rats (N = 7; n = 25).

### Image Data Analysis

X and y movement artefacts were corrected (Template-Matching plug-in for ImageJ) and frames with z movement artefacts removed. Movement shifting was calculated for the SR101 channel and used to correct the OGB-1 image. Neurons were outlined (ROI Manager plug-in, ImageJ, NHI) and fluorescence intensities data for each neuron was processed with a custom software written in MATLAB (Calsignal) (37). ΔF/F0 was computed for each cell, with F0 the mean fluorescence intensity of the recording. Calcium transients were automatically detected (threshold 12 % above baseline, total duration > 1 sec) and then visually examined. For each recording, onsets of calcium transients of each cell were compiled into a binary matrix (38). Pairwise neuronal correlations between cell activities were computed using the normalized cross-correlation for ± 2 frames lag (∼ ± 500 ms). For each pair, 1000 surrogates were generated with the shuffling of calcium onsets while keeping inter-event intervals. The maximum normalized cross-correlation was calculated for each surrogate. A pair of neurons was considered statistically correlated if its true maximum cross-correlation value was not in the 95 % confidence interval computed from surrogates. To detect clusters of neurons, we used the functional clustering algorithm (FCA) (23, 24). This algorithm has no a priori about the number of clusters and groups neurons based on their statistical similitude in temporal patterns. We chose the maximum cross-correlation values as pairwise similarity metrics to group neurons with the FCA and we used surrogates to calculate 95% confidence intervals for each pairwise similarity (24). FCA first merged the activities of the two neurons with the highest maximum cross-correlation value to form a new spike train of activity. This one was then added to the data set from which the two neurons had been removed. This allowed assessing the similarity between the existing cluster and the other neuron activities (24). This procedure was repeated until all active neurons were integrated and as long pairwise similarity was statistically significant (24). We considered as a functional cluster a group of ≥ 3 neurons.

### Statistics

Everywhere, N is the number of rats and n the number of recordings. Statistical tests are anova performed on ranks with cells or recordings nested into rats. A Šidák correction was applied for rejecting the null hypothesis (p < 0.017) when the data was from LSPS synaptic input maps. In Fig. 3B,E we applied the boostrap method and 95 % confidence interval to test a significant difference. Bar graphs and histograms show median values and 25-75^th^ percentiles computed across animals. Exceptions are in Fig. 3B,D,E, Fig. 4E where they show median values and percentiles computed across cells. Symbols in plots of data as a function of postnatal days are each the median value of data from a single rat.

## Acknowledgments

This work was supported by funding from the Institut National de la Santé et de la Recherche Médicale, a grant from the Agence Nationale de la Recherche (SoAbsence, #16-CE37-0021 2016) and a fellowship from the Fondation Française pour la Recherche sur l’Epilepsie (G. Jarre). We thank all the staff of the INMED and GIN animal facilities for providing animal care. We also thank Elodie Fino, Séverine Mahon, Alfonso Represa, and Florian Studer for their critical reading of the manuscript.

## References

1. Schubert D, Kotter R, & Staiger JF (2007) Mapping functional connectivity in barrel-related columns reveals layer- and cell type-specific microcircuits. Brain Struct Funct 212(2):107–119.

2. Shepherd GMG & Svoboda K (2005) Laminar and columnar organization of ascending excitatory projections to layer 2/3 pyramidal neurons in rat barrel cortex. Journal of Neuroscience 25(24):5670–5679.

3. Anastasiades PG, et al. (2016) GABAergic interneurons form transient layer-specific circuits in early postnatal neocortex. Nat Commun 7:10584.

4. Meng X, et al. (2020) Transient Subgranular Hyperconnectivity to L2/3 and Enhanced Pairwise Correlations During the Critical Period in the Mouse Auditory Cortex. Cereb Cortex 30(3):1914–1930.

5. Xue B, Meng X, Xu Y, Kao JPY, & Kanold PO (2022) Transient coupling between infragranular and subplate layers to Layer 1 neurons before ear opening and throughout the critical period depends on peripheral activity. J Neurosci 42(9):1702–1718.

6. Erzurumlu RS & Gaspar P (2012) Development and critical period plasticity of the barrel cortex. Eur J Neurosci 35(10):1540–1553.

7. Fox K (1992) A Critical Period for Experience-Dependent Synaptic Plasticity in Rat Barrel Cortex. Journal of Neuroscience 12(5):1826–1838.

8. Panayiotopoulos CP (1999) Typical absence seizures and their treatment. Arch Dis Child 81(4):351–355.

9. David O, et al. (2008) Identifying neural drivers with functional MRI: an electrophysiological validation. PLoS Biol 6(12):2683–2697.

10. Polack PO, et al. (2007) Deep layer somatosensory cortical neurons initiate spike-and-wave discharges in a genetic model of absence seizures. J Neurosci 27(24):6590–6599.

11. Studer F, et al. (2019) Sensory coding is impaired in rat absence epilepsy. J Physiol 597(3):951–966.

12. Jarre G, et al. (2017) Building Up Absence Seizures in the Somatosensory Cortex: From Network to Cellular Epileptogenic Processes. Cereb Cortex 27(9):4607–4623.

13. Laghouati E, Studer F, Depaulis A, & Guillemain I (2021) Early alterations of the neuronal network processing whisker-related sensory signal during absence epileptogenesis. Epilepsia.

14. Bureau I, Shepherd GM, & Svoboda K (2004) Precise development of functional and anatomical columns in the neocortex. Neuron 42(5):789–801.

15. Shepherd GMG, Pologruto TA, & Svoboda K (2003) Circuit analysis of experience-dependent plasticity in the developing rat barrel cortex. Neuron 38(2):277–289.

16. Finnerty GT & Connors BW (2000) Sensory deprivation without competition yields modest alterations of short-term synaptic dynamics. Proceedings of the National Academy of Sciences of the United States of America 97(23):12864–12868.

17. Bender KJ, Rangel J, & Feldman DE (2003) Development of columnar topography in the excitatory layer 4 to layer 2/3 projection in rat barrel cortex. Journal of Neuroscience 23(25):8759–8770.

18. Marques-Smith A, et al. (2016) A Transient Translaminar GABAergic Interneuron Circuit Connects Thalamocortical Recipient Layers in Neonatal Somatosensory Cortex. Neuron 89(3):536–549.

19. Bureau I, Shepherd GM, & Svoboda K (2008) Circuit and plasticity defects in the developing somatosensory cortex of FMR1 knock-out mice. J Neurosci 28(20):5178–5188.

20. Brill J & Huguenard JR (2009) Robust short-latency perisomatic inhibition onto neocortical pyramidal cells detected by laser-scanning photostimulation. J Neurosci 29(23):7413–7423.

21. Golshani P, et al. (2009) Internally mediated developmental desynchronization of neocortical network activity. J Neurosci 29(35):10890–10899.

22. Martini FJ, Guillamon-Vivancos T, Moreno-Juan V, Valdeolmillos M, & Lopez-Bendito G (2021) Spontaneous activity in developing thalamic and cortical sensory networks. Neuron 109(16):2519–2534.

23. Feldt Muldoon S, Soltesz I, & Cossart R (2013) Spatially clustered neuronal assemblies comprise the microstructure of synchrony in chronically epileptic networks. Proc Natl Acad Sci U S A 110(9):3567–3572.

24. Feldt S, Waddell J, Hetrick VL, Berke JD, & Zochowski M (2009) Functional clustering algorithm for the analysis of dynamic network data. Phys Rev E Stat Nonlin Soft Matter Phys 79(5 Pt 2):056104.

25. De Leon Reyes NS, et al. (2019) Transient callosal projections of L4 neurons are eliminated for the acquisition of local connectivity. Nat Commun 10(1):4549.

26. Chattopadhyaya B, et al. (2004) Experience and activity-dependent maturation of perisomatic GABAergic innervation in primary visual cortex during a postnatal critical period. J Neurosci 24(43):9598–9611.

27. Modol L, et al. (2020) Assemblies of Perisomatic GABAergic Neurons in the Developing Barrel Cortex. Neuron 105(1):93–105 e104.

28. Gainey MA, Aman JW, & Feldman DE (2018) Rapid Disinhibition by Adjustment of PV Intrinsic Excitability during Whisker Map Plasticity in Mouse S1. J Neurosci 38(20):4749–4761.

29. Lambo ME & Turrigiano GG (2013) Synaptic and intrinsic homeostatic mechanisms cooperate to increase L2/3 pyramidal neuron excitability during a late phase of critical period plasticity. J Neurosci 33(20):8810–8819.

30. Studer F, et al. (2022) Aberrant neuronal connectivity in the cortex drives generation of seizures in rat absence epilepsy. Brain.

31. Goncalves JT, Anstey JE, Golshani P, & Portera-Cailliau C (2013) Circuit level defects in the developing neocortex of Fragile X mice. Nat Neurosci 16(7):903–909.

32. Domanski APF, Booker SA, Wyllie DJA, Isaac JTR, & Kind PC (2019) Cellular and synaptic phenotypes lead to disrupted information processing in Fmr1-KO mouse layer 4 barrel cortex. Nat Commun 10(1):4814.

33. Gibson JR, Bartley AF, Hays SA, & Huber KM (2008) Imbalance of neocortical excitation and inhibition and altered UP states reflect network hyperexcitability in the mouse model of fragile X syndrome. J Neurophysiol 100(5):2615–2626.

34. Consortium DBFX (1994) Fmr1 knockout mice: a model to study fragile X mental retardation. Cell 78:23–33.

35. Plantier V, et al. (2018) Direct and Collateral Alterations of Functional Cortical Circuits in a Rat Model of Subcortical Band Heterotopia. Cereb Cortex 29(10):4253–4262.

36. Stosiek C, Garaschuk O, Holthoff K, & Konnerth A (2003) In vivo two-photon calcium imaging of neuronal networks. Proc Natl Acad Sci U S A 100(12):7319–7324.

37. Platel JC, et al. (2005) Na+ channel-mediated Ca2+ entry leads to glutamate secretion in mouse neocortical preplate. Proc Natl Acad Sci U S A 102(52):19174–19179.

38. Cossart R, Aronov D, & Yuste R (2003) Attractor dynamics of network UP states in the neocortex. Nature 423(6937):283–288.

